# Integrative and conjugative elements and their hosts: composition, distribution, and organization

**DOI:** 10.1101/135814

**Authors:** Jean Cury, Marie Touchon, Eduardo P. C. Rocha

## Abstract

Conjugation of single-stranded DNA drives horizontal gene transfer between bacteria and was widely studied in conjugative plasmids. The organization and function of integrative and conjugative elements (ICE), even if they are more abundant, was only studied in a few model systems. Comparative genomics of ICE has been precluded by the difficulty in finding and delimiting these elements. Here, we present the results of a method that circumvents these problems by requiring only the identification of the conjugation genes and the species’ pan-genome. We delimited 200 ICEs and this allowed the first large-scale characterization of these elements. We quantified the presence in ICEs of a wide set of functions associated with the biology of mobile genetic elements, including some that are typically associated with plasmids, such as partition and replication. Protein sequence similarity networks and phylogenetic analyses show that ICEs are modular and that their gene repertoires can be grouped in function of their conjugation types, even if integrases were shown to be paraphyletic relative to the latter. We show that there are general trends in the functional organization of genes within ICEs and of ICEs within the bacterial chromosome paving the way for future functional and evolutionary analyses.

## Introduction

Bacterial diversification occurs rapidly by the constant influx of exogenous DNA by horizontal gene transfer (HGT) (1-3). As a consequence, the diversity of genes found in the strains of a species, its pangenome, is usually much higher than the number of genes found in a single bacterial genome at a given time (4). The pangenome represents a huge reservoir of potentially adaptive genes, whose potential has become evident in the rapid spread of antibiotic resistance in the last decades (5), and in the emergence of novel pathogens (6). Mobile genetic elements (MGE) drive the spread of genes in populations using a variety of mechanisms, often encoded by the elements themselves.

Conjugative MGEs carry between a few dozens to several hundred genes (7). They can be extra-chromosomal (plasmids) or integrative (ICEs). Conjugation requires an initial step of cell-to-cell contact during mating pair formation (MPF). The mechanism is the same for plasmids and ICEs, apart from the initial and final steps of chromosomal excision and integration (8,9). The mechanism of transfer proceeds in three steps. Initially, the relaxase (MOB) nicks the DNA at the origin of transfer (*oriT*), and binds covalently to one of the DNA strands. The nucleoprotein filament is then coupled to the type 4 secretion system (T4SS) and transferred to the recipient cell. Finally, the element is replicated in the original and novel hosts leading to double stranded DNA molecules in each cell. One should note that some integrative elements are transferred using another mechanism, also called conjugation, relying on double-stranded DNA. They are restricted to certain Actinobacteria, have been recently described in detail (10,11), and will not be mentioned in this work. Integration of ICEs is usually mediated by a Tyrosine recombinase (integrase), but some ICEs use Serine or DDE recombinases instead (9,12-14). There are eight types of MPF(15,16), each based on a model system as described in Figure 1. The identification of a valid conjugative system requires the presence of a relaxase, of the VirB4 ATPase, a coupling ATPase (T4CP), and several other proteins that may differ between types (although they are sometimes distant homologs or structural analogs) (16). Both ICEs and plasmids can be of any MPF type, but some preferential associations have been observed: there are few plasmids of MPF_G_ and few ICEs of MPF_I_ and MPF_F_ (17).

**Figure 1:**
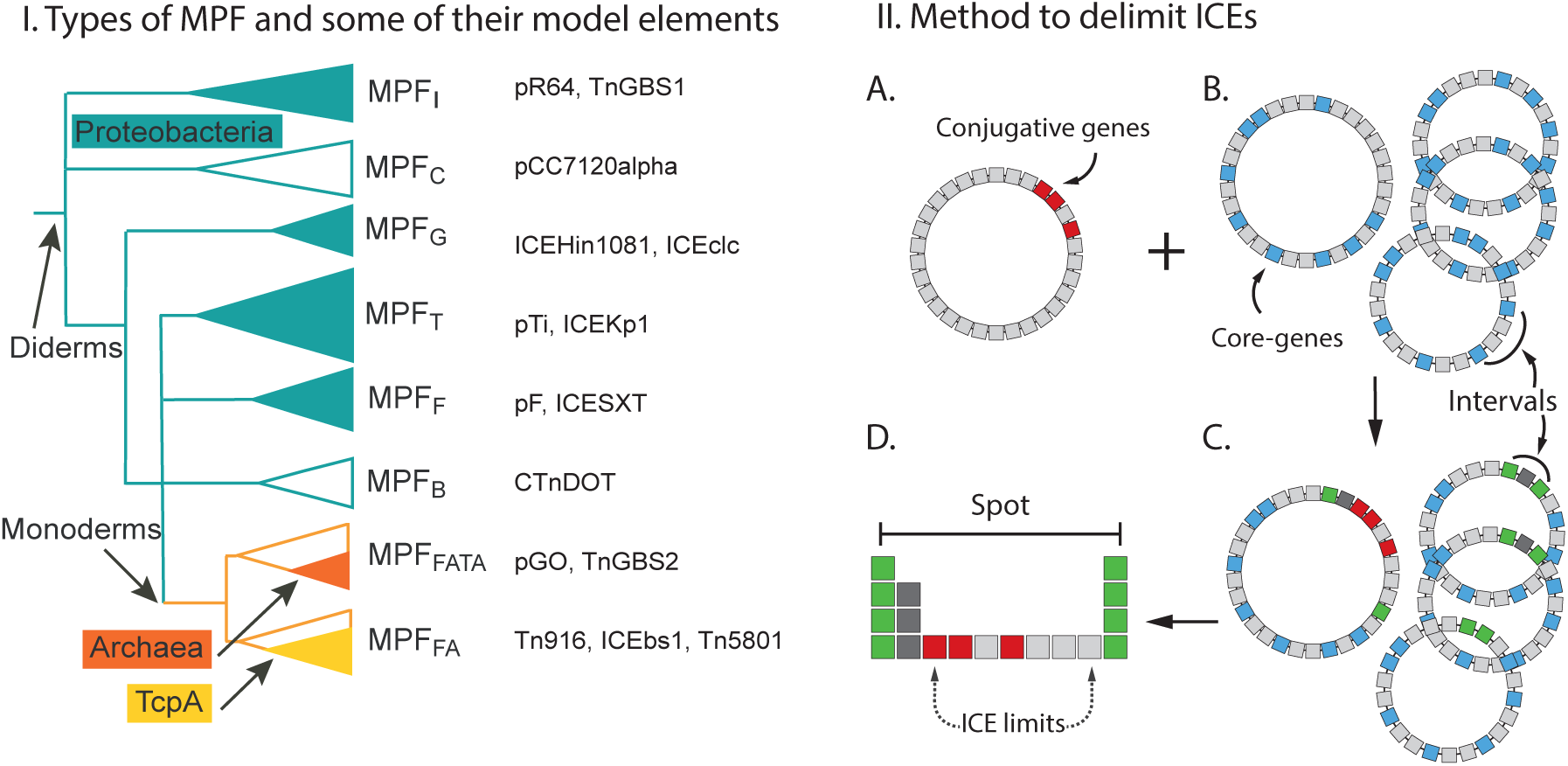
Scheme of the MPF types and procedure for ICE delimitation. **I.** The phylogenetic tree displays the evolutionary relationships between MPF types as given by the VirB4 phylogeny. Most lineages are from diderm (green branches), the systems from monoderms (yellow branches, including Firmicutes, Actinobacteria, Archaea, and Tenericutes) being derived from these. MPF_B_ (Bacteroides) and MPF_C_ (Cyanobacteria) were absent from our data because of lack of enough genomes for the analysis. The full green clades indicate systems that are typically found in Proteobacteria. The label TcpA indicates a clade that used this protein as T4CP (an homologous ATPase from the typical T4CP). In front of each tip of the tree, there is a non-exhaustive list of well-known conjugative elements (ICE or plasmid) for each MPF type. The tree was taken from (15). **II. A.** Genes encoding conjugative systems (Red) were detected in bacterial genomes using MacSyFinder. **B**. We restricted the dataset of ICEs to those present in the 37 species for which we had at least four genomes and an ICE. We built the core genome (blue, core-genes) of each species. An interval is defined as the region between two consecutive core-genes. **C**. The information on the conjugative system and the core-genes is used to delimit the chromosomal interval harboring the ICE. An ICE is thus flanked by two core genes that provide an upper bound for its limits (in green). **D**. The two core genes define intervals in a several genomes of the species (typically in all of them), and the set of such intervals is called a spot. We built the spot pan-genome, *i.e.*, we identified the gene families present in the spot, and mapped this information on the interval with the conjugative system. Finally, the manual delimitation is based on a visual representation of the spot including this information and G+C content (see Figure S1 and Methods).

ICEs are more numerous than conjugative plasmids among sequenced genomes (17), but their study is still in the infancy. Beyond the fact that they were discovered more recently, the extremities of ICEs are difficult to delimit precisely in genomic data. Hence, most data available on the biology of ICEs comes from a small number of experimental models, such as the ICE SXT of the MPF_F_ type, ICEclc (MPF_G_), MlSym^R7A^ (MPF_T_), Tn916 and ICEBs1 (MPF_FA_), CTnDOT (MPF_B_) (18-21). Several of these elements encode traits associated with pathogenicity, mutualism, or the spread of antibiotic resistance, which spurred the initial interest on the ICE. It has been suggested that this has biased the study of ICE biology and that analyses with fewer *a priori* are needed to appreciate the evolutionary relevance of these elements (9). Some ICEs encode mechanisms typically found in plasmids, such as replication and partition (22-25), and phylogenetic studies showed that interconversions between ICE and conjugative plasmids were frequent in the evolutionary history of conjugation (17). Together, these results suggest that ICE and CP are more similar than previously thought (26). Interestingly, ICEs also encode functions typically associated with other MGEs, such as phage-related recombinases (27), and transposable elements (28).

The unbiased study of the gene repertoires and structural traits of ICEs is important to improve the current knowledge on these elements. Here, we developed a method based on the comparison of multiple genomes in a species to identify, class, and study ICEs in bacteria. The method allowed us to delimit ICEs, analyze their gene content, study the resemblance between elements and their internal organization, and to study their distribution in chromosomes. Our methodology does not use any *a priori* knowledge on the organization of previously known ICEs, apart from requiring the presence of a conjugative system. Hence, it should not be affected by ascertainment biases caused by the use of model systems to identify novel ICEs. We show that it provides a broad view of the diversity of ICEs among bacterial genomes.

## Material and Methods

### Data

The complete genomes of 2484 Bacteria were downloaded from NCBI RefSeq (http://ftp.ncbi.nih.gov/genomes/refseq/bacteria/), in November 2013. We used the classification of replicons in plasmids and chromosomes as provided in the GenBank files. We searched for conjugation systems in all replicons of all genomes. Yet, the delimitation of ICEs was restricted to species having at least one chromosomally encoded conjugative system and at least four genomes completely sequenced (37 species, 506 genomes). Sequences from experimentally validated ICEs were retrieved form the ICEberg database version 1.0 (http://db-mml.sjtu.edu.cn/ICEberg/).

### Detection of conjugative systems

Conjugative systems were found with MacSyFinder (29), using protein profiles and definitions following the previous work of (16) (File S1). An element was considered as conjugative when it contained the following components of the conjugative system: VirB4/TraU, a relaxase, a T4CP, and a minimum number of MPF type-specific genes: two for types MPF_FA_ and MPF_FATA_, or three for the others. MacSyFinder was ran independently for each given MPF type with default parameters (hmmer e-value < 0.001, protein profile coverage in the alignment higher than 50%). Conjugative elements of some taxa lack known relaxases, *e.g.*, some Tenericutes and some Archaea. Since T4SS can be mistaken by protein secretion systems in the absence of relaxases, such systems were excluded from the analysis. We have set up a CONJscan module for MacSyFinder (downloadable at https://github.com/gem-pasteur/Macsyfinder_models) to identify these elements using a Unix-like command line. The models in CONJscan can be modified by the user. We also setup a Webserver (https://galaxy.pasteur.fr/) to identify these elements.

### Identification of gene families, core and pan-genomes, spots, and intervals

We defined within each species a core genome, a set of intervals, and a set of spots. The core genome is the set of families of orthologous proteins present in all genomes of the species. We computed the core genome of each species as in (30). Briefly, orthologous genes were identified as the bi-directional best hits (BBH, using global end-gap-free alignments with more than 80% of protein similarity, less than 20% of difference in length, and having at least four other pairs of BBH hits within a neighborhood of ten genes), and the core genome was defined as the intersection of the pairwise lists of orthologues between genomes using the reference strain as a pivot.

We defined intervals as the regions between two consecutive core genes in a genome. We defined a spot as the set of intervals flanked by members of the same pair of core gene families (Figure 1). Hence, a spot has either one interval per genome or none (if the interval was split by a rearrangement). If the region has not endured chromosomal rearrangements (the most typical situation), then the spot will contain as many intervals as the number of genomes. We identified the gene families of the spots that encode at least one ICE (spot pan-genome). For this, we searched for sequence similarity between all proteins in the spot using blastp (version 2.2.15, default parameters) and used the output to build protein families using Silix (version 1.2.8) (31). Proteins whose alignments had more than 80% identity and at least 80% of coverage were grouped in the same family. The members of the spot pan-genome that were not part of the core genome constituted the accessory genome.

### Delimitation of ICEs

We analyzed conjugative systems encoded in chromosomes, and delimited the corresponding ICEs using comparative genomics. There is usually a high turnover of ICEs at the species level (*i.e.*, most elements are present in only a few strains), implicating that a few genomes are usually sufficient to delimit the element by analyzing the patterns of gene presence and absence. We restricted our analysis to species with at least four genomes completely sequenced and assembled (without gaps). ICEs were delimited in two steps. First, we identified the spots encoding conjugative systems (see definition above). The core genes flanking these spots provide upper bounds for the limits of the ICE. Second, we analyzed the interval with the ICE and identified the limits of the element by overlaying the information on the presence of genes of the conjugation system, on G+C content, and on the frequency of accessory genes in the spot pan-genome. The genes of the ICE are expected to be present in the spot pan-genome at similar frequencies (some differences may be caused by mutations, deletions, transposable elements, and annotation errors) and this information is usually sufficient to delimit the ICE. We produced a visual representation of this data in the context of the spot, and used it to precisely delimit the ICE at the gene-level (see Figure 1 and Figure S1).

### Specific functional analyses

Antibiotic resistance genes were annotated with the Resfams profiles (core version, v1.1) (32) using HMMER 3.1b1 (33), with option --cut_ga. The cellular localization was determined with PsortB (version 3.0) (34), using the default parameters for diderms and monoderms separately. Genes encoding stable RNAs were annotated using Infernal (35) and Rfam covariance models (36) (hits were regarded as significant when e-value ≤ 10^-5^). Integrons were detected using IntegronFinder v1.5 with the --local_max option (37). DDE transposase were annotated with MacSyFinder (29) following the procedure described in (30). Integrases were detected with the PFAM profile PF00589 for tyrosine recombinases and the pair PF00239 and PF07508 for Serine recombinases (http://pfam.xfam.org/) (38). HMMER hits were regarded as significant when their e-value was smaller than 10^-3^ and their alignment covered at least 50% of the protein profile.

Specific HMM protein profiles were built with HMMER v3.1b1 for partition systems (39,40), replication proteins (41), and entry exclusion systems (42). In the general case, we started from a few proteins with experimental evidence of the given function, curated by experts, or reported in published databases. Since these sets were usually small and present in a small number of species, we used a two-step procedure (described below): we started by building preliminary profiles, used them to scan the complete genome database, and then used these results to make the final profiles. First, the proteins of experimental model systems were aligned with mafft v7.154b (with --auto parameter) (43), and manually trimmed at the N-and C-terminal ends with SeaView v.4.4.1 (44).

The alignments were used to make preliminary HMM profiles using hmmbuild from HMMER v.3.1b1. A first round of searches with these profiles using hmmsearch (e-value < 10^-3^ and coverage >50%) returned hits that were clustered with usearch (--cluster_fast at 90% identity) (45). We took only the longest protein of each cluster, to remove redundant sequences, and searched for sequence identity between all pairs of these representative hits using blastp v.2.2.15 (with the –F F parameter to not filter query sequences). The result was clustered with Silix (version 1.2.8, 40% identity and 80% of coverage). We made multiple alignments of the resulting families and used them to build a novel set of HMM protein profiles (alignment and trimming as above). The detailed procedure used for the protein profiles of each function is given in the supplementary material.

### Functional annotation

We used HMMER v.3.1b1 (e-value<0.001 and coverage of 50%) to search ICEs for hits against the EggNOG Database of hmm profiles (Version 4.5, bactNOG). These results were used to class genes in broad functional categories. We added a class “Unknown” for genes lacking hits when queried with EggNOG. We tested the over-representation of given functional categories in ICE using binomial tests where the successes were given by the number of times that the relative frequency of a given functional category was higher in the ICE than in the rest of the host chromosome. Under the null hypothesis, any given category is as frequent in the ICE as in the rest of the chromosome (relative to the number of genes). Formally:

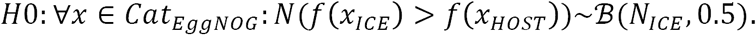

With *N*(*f x*_*ICE*_) > *f*(*x*_*HOST*_) being the number of times the EggNOG category ***x*** had a higher relative frequency in the ICE than in the rest of the host chromosome, and with *B*(*N*_*ICE*_, 0.5) a binomial distribution with *N*_*ICE*_ successes (199 typed ICEs) and an *a priori* probability of 50%.

### Networks of homology

We searched for sequence similarity between all proteins of all ICE elements using blastp v.2.2.15 (default parameters) and kept all bi-directional best hits with an e-value lower than 10^-5^. We used the results to compute a score of gene repertoire relatedness for each pair of ICEs weighted by sequence identity:

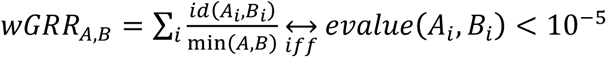

where (*A*_*i*_, *B*_*i*_) is the pair *i* of homologous proteins in ICEs A and B, id(A_i_,B_i_) is the percentage of identity of their alignment, min(A,B) is the number of proteins of the element with fewest proteins (A or B). The wGRR represents the sum of the percentage of identity for all pairs of homologous proteins between ICE A and B, divided by the number of proteins of the smallest ICE. A similar score was previously used to compare prophages (46). We kept pairs of ICEs with a wGRR higher than a certain threshold (5% for the general analysis and 30% for the supplementary analysis). A wGRR of 5% represents, for instance, the occurrence of one homologous gene with 100% identity between two ICEs, where the smallest encodes 20 proteins. We computed two versions of this score for each pair of ICEs, one with all proteins and another after excluding the proteins from the conjugative system.

### Phylogenetic tree

The phylogenetic analysis used the 134 integrases from ICEs containing exactly one tyrosine recombinase. We excluded the other elements because additional recombinases might be involved in other functions (dimmer resolution, DNA inversions) and could confound the results. We added to this dataset: 60 phage integrases randomly selected from a database of 296 non-redundant (<90% identity) phage integrases from RefSeq; 11 integrases from pathogenicity islands (Table 2S from (47)); 25 experimentally studied XerC and XerD from (48), XerS from (49), XerH from (50,51); seven integrases representing the diversity of integron integrases (37). The final dataset was composed of 237 integrases. The initial alignment was made with mafft (with parameters --maxiterate 1000 --genafpair), and 100 alternative guide-trees were built.

We then removed columns with less than 50% of confidence score, as calculated by GUIDANCE 2 (52). We used Phylobayes MPI (version 1.7, model CAT+GTR+Gamma, two chains for 30 000 iterations) to build the phylogeny from that alignment (53). Following the guidelines of Phylobayes, we considered that chains had converged when the maximum difference across all bipartitions was lower or equal than 0.3 (0.24). The tree was represented with Figtree v1.4.2.

### Identification of origin and terminus of replication

The origin (ori) and terminus (ter) of replication were those predicted in DoriC (54). When the ratio of the predicted replichores length was greater than 1.2, the genome was removed from the corresponding analysis. The leading strand was defined as the one showing a positive GC skew (55).

## Results and Discussion

### Identification and delimitation of ICEs by comparative genomics

We developed a procedure to identify and delimit ICEs in two stages (see Materials and Methods). First, we identified conjugative systems in bacterial chromosomes using our published methodology, previously shown to be highly accurate (17,56). We then concentrated our attention on the conjugative systems of species for which at least four complete genomes were available. We defined for each species the core and pangenomes, the set of intervals (locations between consecutive core genes), and their spots (set of intervals flanked by the same families of core genes, see Methods). Each ICE was then placed on the corresponding interval and delimited based on the analysis of the frequency of gene families in the spot pan-genome, the position of conjugative systems, and the G+C content (Figure 1 and S1).

We identified five pairs of ICEs in tandem. They were relatively easy to identify with our procedure because the corresponding intervals had two copies of the conjugation apparatus (and typically two integrases). The tandem ICEs corresponded to different MPF types in four out of five pairs. This fits previous observations that tandems of identical ICEs are very rare in *recA*^+^ backgrounds (all the ICEs analyzed in this work are in these circumstances) (57). Some elements encoded two or more integrases or relaxases and only one conjugation system. They may be composite elements resulting from the independent integration of different elements, or they may correspond to ICE encoding additional tyrosine recombinases with other functions (than that of being an integrase). When faced with such elements, and when the frequency of the genes was homogeneous, we decided to include them in one single ICE. This choice was based on the published works showing that ICEs can mobilize composite elements, including genomic islands or IMEs (58-60), and on the observation that conjugative plasmids sometimes also include other integrative elements and multiple relaxases (7).

Of note, we were not able to obtain a general method to identify accurately the attachment sites (*attL* and *attR)* delimiting ICEs. This also complicates the separation between simple and composite ICEs. Hence, we placed the elements’ borders at the position of their flanking genes. The curation process started with 601 conjugative systems detected among 2484 complete genomes. We selected 207 (∼35%) of these because they were present in species with more than four complete sequenced genomes. Within these, we were able to delimit 200 ICEs within 37 species. Among them, 118 ICEs were present in 41 spots showing that integration of different ICEs in the same region is common. To assess if ICEs in the same spot are different elements or the result of single ancestral integrations in the genome, we computed the weighted gene repertoire relatedness (wGRR), which represents the proportion of homologous genes between two ICEs weighted by their sequence identity (see Methods, Figure S2). Most ICEs had very low wGRR with the others. Yet, there were 61 (70) ICEs with wGRR>90 (wGRR>80) with at least one other ICE among those in the same spot. If one had kept only one ICE per family with this threshold of wGRR>90, this would have reduced the number of ICEs from 200 to 160 (Figure S2). Yet, these elements are not necessarily derived from the same ancestral event of integration since we found 29 ICEs with such high wGRR values in different species and 24 in different genera. Hence, instead of arbitrarily select one ICE per spot because they can be the result of a common ancestral integration event, we opted to maintain all elements in the subsequent analyses and control for this effect, when necessary.

We used information from the ICEberg database to validate the delimitation procedure. We could not use the whole database because there was no complete conjugative system in 51% of the elements of ICEberg (184 out of 358), and 40% of them even lacked the essential protein VirB4/TraU. Hence, we restricted our comparisons to the 19 ICEs in ICEberg that were derived from experimental data or predicted from literature and that were in a genome available in our dataset (Table S1). Among these elements, 16 ICEs had their start and end positions within less than 2.5 kb of ours (typically less than 500bp, Figure S3). We analyzed in detail the three cases showing discordance between the two datasets. Two of them corresponded to a tandem of ICEs that we split in our procedure because they had two different complete conjugative systems. They were identified as a single ICE in ICEberg. The third problem arose because the ICEberg annotation included an MPF_FATA_ ICE interrupted by an MPF_FA_ ICE. Our procedure spotted the complete MPF_FA_ ICE. These results suggest that our delimitation is more accurate. It is important to note that our procedure does not use any information on the experimental models of ICE; it is only based on the identification of conjugation proteins, and the frequency of accessory genes. Hence, the accuracy of our method is expected to remain high even when studying poorly known ICE families. It is also expected to improve upon inclusion of more genomes within a species.

### General features of ICEs

We classed the 200 delimited ICEs in function of the MPF and relaxase (MOB) types. We identified five of the eight previously defined MPF types among these elements: type F (9), FA (72), FATA (29), T (49), and G (40). Three types were absent from our data because they are strictly associated with clades for which not enough complete genomes were available (MPF_B_ for Bacteroides and MPF_C_ for cyanobacteria) or corresponded to systems that were previously shown to be extremely rare among ICE (MPF_I_) (17). Accordingly, the distribution of MPF types differed from that of the 601 conjugative systems identified in all chromosomes (χ^2^, p-value<0.001, Figure S4), and included mostly Firmicutes (for MPF_FA_ and MPF_FATA_) and Proteobacteria (for the others). One ICE had an undetermined MPF type (albeit it included a VirB4, a MOB and a coupling protein), and was excluded from the remaining analyses (except from the wGRR network, see below). Expectedly, the types of relaxases identified in the ICEs corresponded to those frequently found in Proteobacteria and Firmicutes (17) (Figure S5).

The size of ICEs in the scientific literature varies between ∼13kb (ICE*Sa1* in *Staphylococcus aureus* (61)) and ∼500kb (ICEMlSymR71 in *Mesorhizobium loti* R7A (62)). In our dataset, the smallest ICE was also identified in *S. aureus* (strain USA300-FPR3757) and was only 11.5 kb long. The largest ICEs were found in *Rhodopseudomonas palustris BisB18* (MPF_T_) *and Pseudomonas putida S16* (MPF_G_) and were around 155 kb long. The distribution of ICE size per MPF type showed that type F (median size of 99 kb) and G (median 80 kb) were the largest, whereas those of type FA were the smallest (median 23.5kb) (Figure 2). The distribution of the size of ICE in our dataset is close to that of ICEs reported in the literature, even if we could not identify any ICE of a size comparable to ICEMlSymR71. Such large ICEs may be rare or specific of taxa not sampled in our study. ICEs were found to be AT-rich compared to their host’s chromosome (Figure 2), which is a general trend for mobile genetic elements (63). The size distribution of 160 families of ICEs is similar to the one for the entire dataset (Figure S6), validating our decision to keep all ICEs in our analysis.

**Figure 2:**
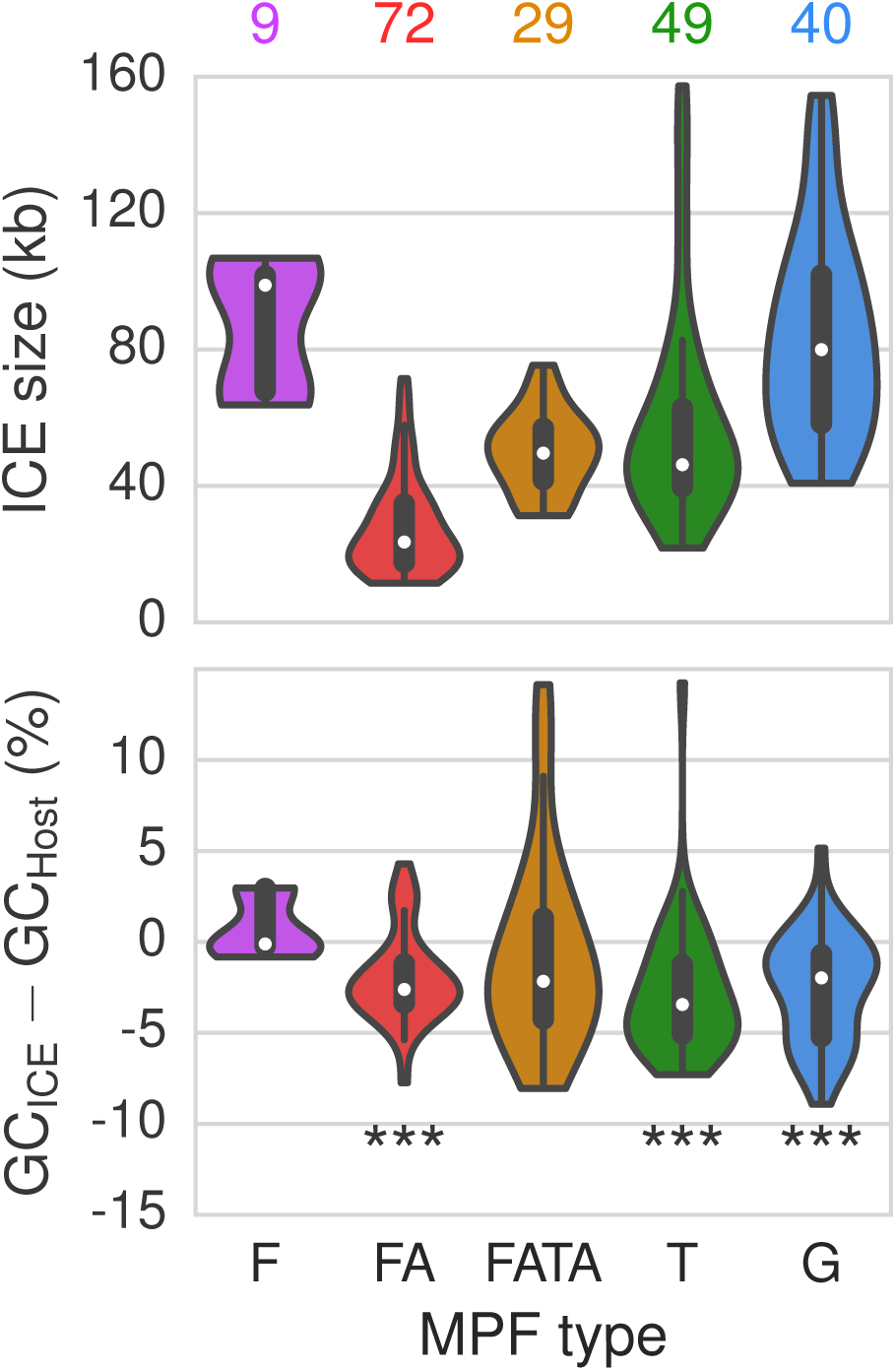
ICE statistics as a function of the MPF type. Top. Distribution of the size of ICEs (in kb). The numbers above each violin plot represent the number of elements in each category. Bottom. Distribution of pairwise differences between the GC content of the ICE and that of its host. The violin plots represent the kernel density estimation of the underlying values. Here the violin plots is limited by the minimum and maximum values. ***: p-value<0.001, Wilcoxon signed-rank test (rejecting the null hypothesis that the difference is equal to zero).

It was known that the largest plasmids are typically in the largest genomes (7). The size of ICE and their host chromosomes are also correlated (ρ=0.47, p-value<0.0001, after discounting the ICE size from chromosome size), even if this is partly because the smaller types of elements are associated with clades with the smallest genomes (Figure S7). Larger ICEs may be disfavored in smaller genomes because they are harder to accommodate in terms of genome organization (64), thus leading to higher fitness cost, or because smaller genomes endure lower rates of HGT (65). Interestingly, the average size of ICEs (52.4 kb) could be a general feature of integrative elements, since it comes near to that of temperate phages (∼50 kb) of enterobacteria (66), and of known pathogenicity islands (ranging between 10kb and 100 kb (67)).

### The families of ICEs

We analyzed the network of protein sequence similarity between ICEs in relation to some of the best-known experimental models (SXT, ICEclc, Tn916, ICEBs1, TnGBS2, ICEKp1). To this end, we used the abovementioned weighted gene repertoire relatedness (wGRR) scores between every pair of ICEs. The network of wGRR-based relationships between ICEs was represented as an undirected graph, where nodes represent ICEs and edges were weighted according to the wGRR (if wGRR>5) (Figure 3). This graph showed that all ICEs were connected in a single component (all nodes can be accessed from any others) with the exception of a group of ICEs from *H. pylori*. ICEs grouped predominantly according to their MPF types, which were sometimes split in several clusters. Interestingly, ICEs in Firmicutes (FA and FATA) and Proteobacteria (T, G and F) formed two higher groups, even if sub-groups typically include different species. This fits the observation that FA and FATA are sister-clades in the phylogeny of VirB4 (15). Interestingly, a similar graph where MPF proteins were excluded from the calculation of wGRR produced very similar results, suggesting that the effect of the host taxa may be preponderant in the split into two groups according to the major phyla (Figure S8). To detail the similarities between ICEs, we also clustered the ICEs using a more restrictive threshold (wGRR > 30, Figure S9). This graph is composed of many connected components of single MPF types, highlighting the huge diversity of ICEs.

**Figure 3:**
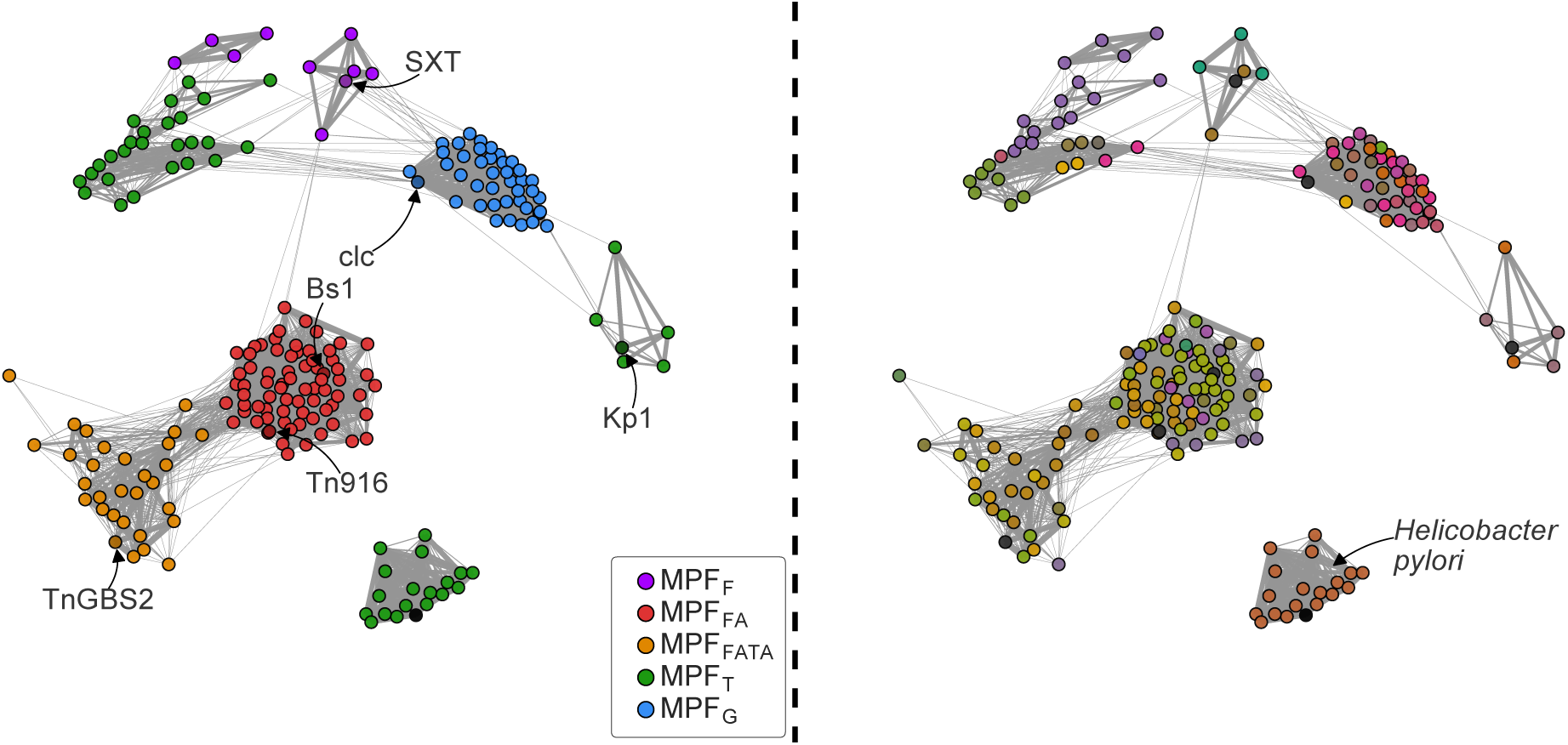
Representation of the wGRR-based network of ICEs. The nodes represent the ICEs and the edges link pairs of ICEs with wGRR score above 5% (the thickness of the edge is proportional to the score). On the left, nodes are colored according to the MPF type. Darker nodes represent ICEs commonly used as experimental models, and are indicated by arrow. On the right, nodes are colored according to the species of the host to highlight the distribution of the 37 species. The information of the species and type are in supplementary tables S4. The position of the point has been determined by the Fruchterman-Reingold force-directed algorithm, as implemented in the NetworkX python library (spring layout).

### Independent integrase acquisitions by ICE

Integrases allow the integration of ICEs in the chromosome and are one of their most distinctive features (relative to conjugative plasmids). Around 70% of the ICEs encoded a single tyrosine recombinase, nine encoded a Serine recombinase, 21 had at least two integrases, among which six had both Serine and Tyrosine recombinases. A total of 37 ICEs lacked integrases, among which six encoded one or more DDE recombinases (in two cases the genes are at the edge of the ICE) and six encoded pseudogenized integrases. DDE recombinases-mediated ICE integration was previously described for ICE TnGBs2 in *Streptococcus agalactiae* (13) and for ICEA in *Mycoplasma agalactiae* (14). Experimental work will be necessary to test if some of the six ICEs with only DDE recombinases use them to replace integrases. Of note, we actually found more transposases in ICEs with Serine or Tyrosine recombinases than in those lacking them (χ^2^ on a contingency table, p-value<0.01). Recently, it has been shown that some ICEs may use relaxases instead of integrases to integrate the chromosome when there is an *oriT* in the genome that can be recognized by the relaxase (68).

A previous analysis using integrases from the tyrosine recombinase family of genomic islands, phages, and six ICEs (of which four of the SXT family), showed that ICE integrases clustered separately (69). However, doubts have been casted on this analysis because of the small number of ICEs used in the analysis (70). We have thus made a phylogenetic tree of tyrosine recombinases from the ICEs encoding one single integrase (Figure 4). We analyzed a total of 237 tyrosine recombinases from ICEs and other elements including integrons, four different types of recombinases involved in chromosome dimer resolution (XerCD, XerS, XerH), pathogenicity islands, and phages (see Methods, Table S2). This tree showed that the Xer recombinases and the integron integrases were all monophyletic. In contrast, genomic islands and ICEs were scattered in the tree. Even ICEs of similar MPF types are systematically paraphyletic. A particularly striking example is provided by a clade in the tree (arc in Figure 4) that contains integrases from several phages, and two types of ICE (Type G and type T) from different species (*K. pneumonia*e and *P. fluorescens*), as well as three different groups of pathogenicity islands. The clear paraphyly of the integrases at the level of MPF types suggests that conjugative elements often exchange the key genes allowing chromosomal integration with otherwise unrelated mobile genetic elements.

**Figure 4:**
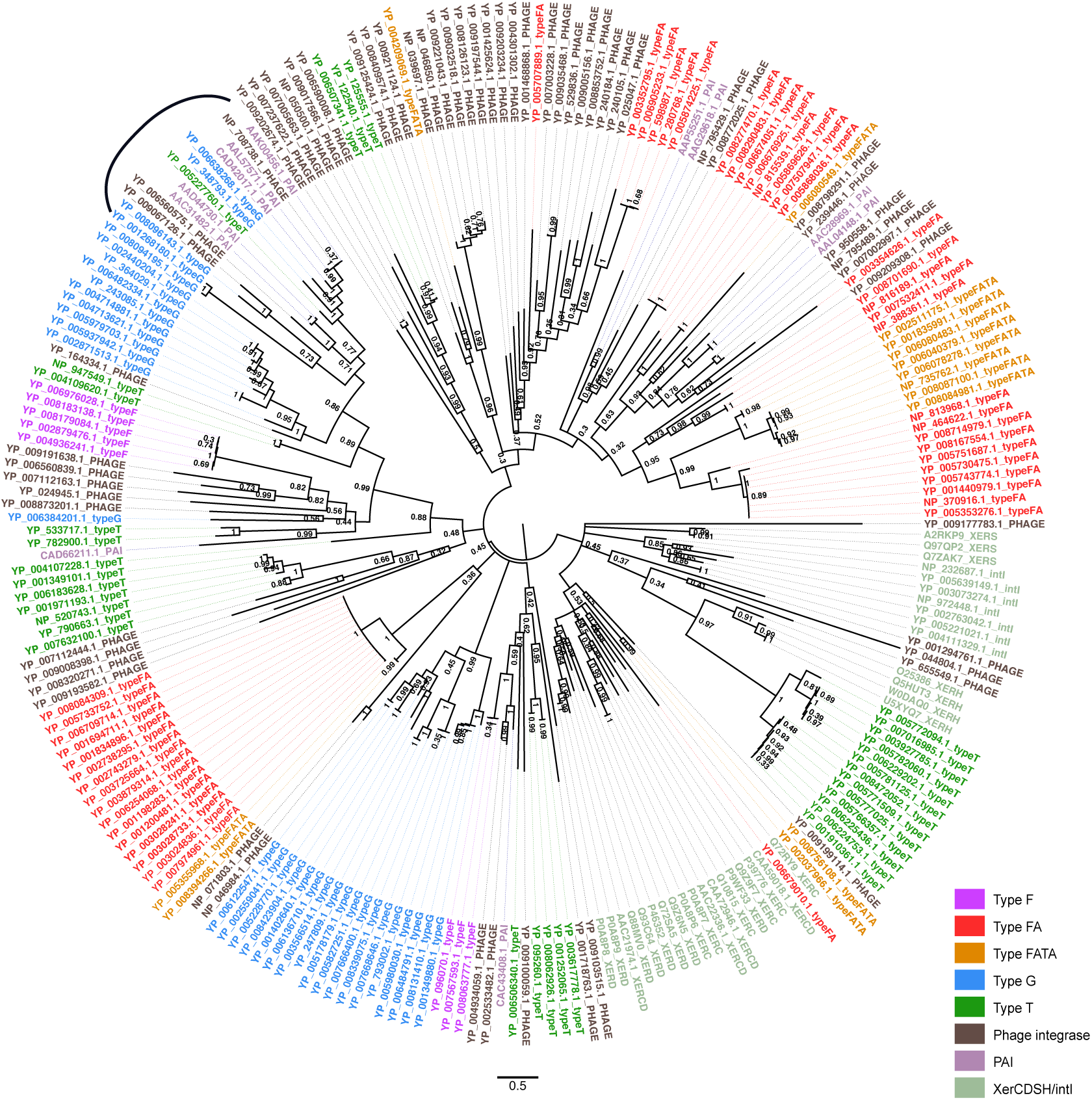
Phylogenetic tree of tyrosine recombinases. The phylogenetic tree was built using 60 prophage integrases (labelled as “…_PHAGE”, black), 11 integrases from pathogenicity islands (“…_PAI”), 25 XerC,D,S or H (“XER”, grey), 7 integron-integrases (“.._intI”, grey), and 134 integrases from ICEs (colored after the MPF type). The tree was built using Phylobayes and the values represent posterior probabilities support of the partition, with a cut off equal to 0.3 (under which nodes are collapsed) (see Methods). The black arc denotes a clade with good support, which contains integrases from prophages, PAI and different MPF types, that is explicitly cited in the text.

### The functional repertoire of ICE

We investigated the functional classification of the genes in ICEs in relation to those in the rest of the host chromosome using the EggNOG database (see Methods). Unknown or unannotated functions accounted for 61% of all genes in ICEs. We observed three functional categories that were systematically more frequent in ICEs (p-value<0.01 with Bonferroni correction for multiple test, Figure 5), including typical ICE functions: secretion (genes related to conjugation), replication/recombination/repair (integrases, relaxases, and transposable elements), and to a lesser extent cell cycle control/cell division/chromosome partitioning. The category associated with transcription (gene expression regulation) showed similar frequencies in the ICE and in the host chromosome. Most functions were systematically less frequent in ICEs. Removing from the analysis the proteins implicated in conjugation did not reveal novel families over-represented in ICEs (Figure S10).

**Figure 5.**
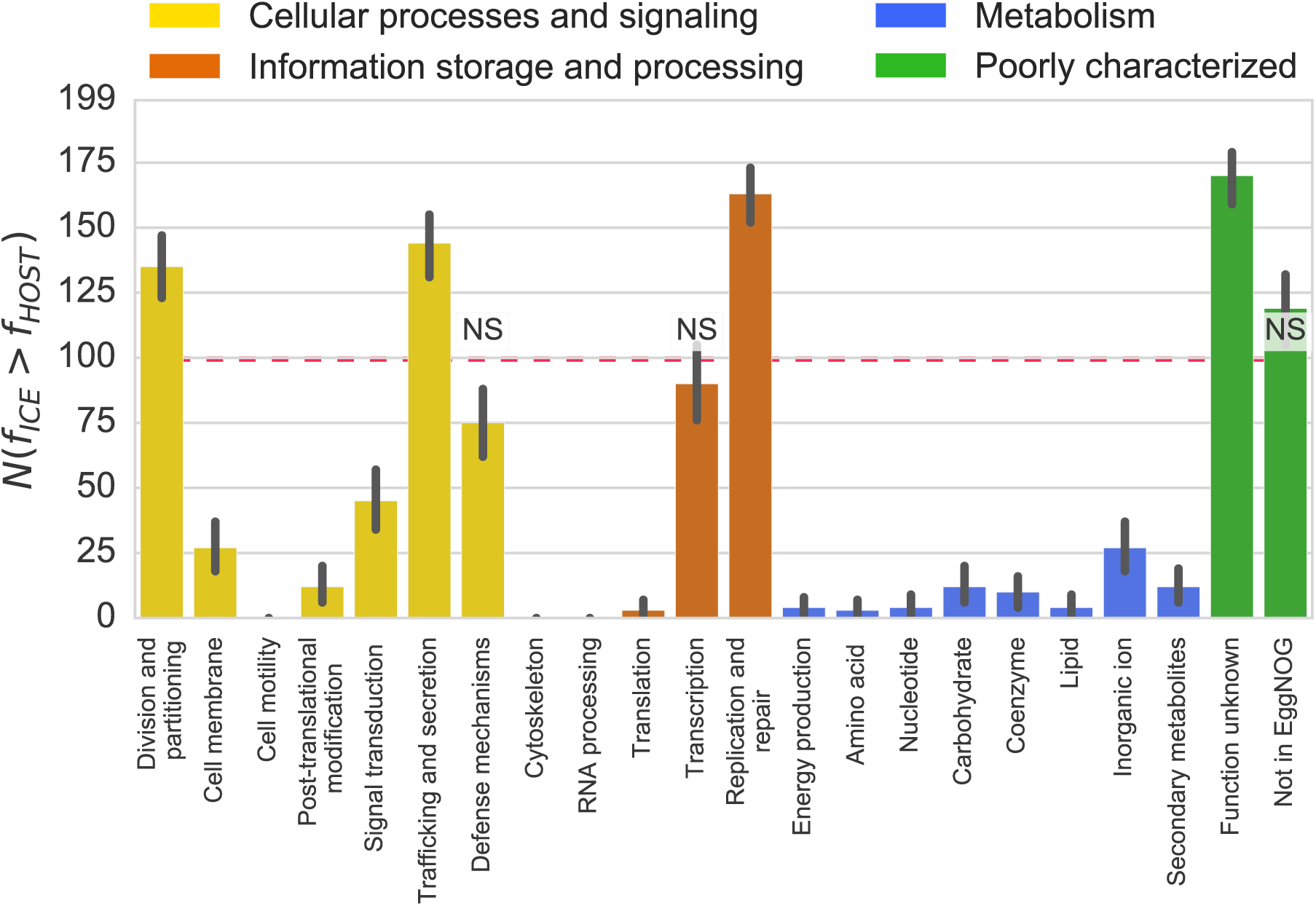
Representation of EggNOG functional categories in ICEs relative to the host chromosome. The bars represent the number of times a given category is found more frequently in an ICE than in its host chromosome (N(f_ICE_>f_HOST_)). The red dotted line represents the expected value under the null hypothesis, where a category is in similar proportion in ICE and its host’s chromosome. Bars marked as NS represent a lack of significant difference (p>0.05, Binomial test with 199 trials and expected value of 0.5), whereas the others are all significantly different (p<0.05, same test). Note that there are 199 trials because one of the 200 ICEs could not be types and was thus excluded, see text. Error bars represent 95% confidence interval computed with 1000 bootstraps. ″Not in EggNOG″ represents the class of genes that didn′t match any EggNOG profile.

Since the functional analysis of ICEs pinpointed an over-representation of functions typical of plasmids, we developed more specific approaches to characterize them (see Methods). We identified 23 partition systems, 13 from type Ia, five of type Ib, and five of type II (no type III). The presence of partition systems suggests the existence of replication systems (even if relaxases can themselves be implicated in ICE replication (25,71)). We found 16 ICEs with proteins predicted to be associated with theta replication (15 in MPF_T_ and one in MPF_G_) and two associated with rolling circle replication (all in MPF_FATA_). Interestingly, all six MPF_T_ ICEs with a partition system also encoded a replication protein. Apart from these traits (MPF type, partition and replication systems), these six ICEs are very different (average wGRR score of 34%). To the best of our knowledge ICE replication has not been reported in this family (the most numerous one among complete genomes). Naturally, if some of the relaxases act as replicases, then the actual number of replicases in ICE could be much larger. This might explain why few or no replication systems were found in type F and G although they encode partition systems.

Several ICEs encode accessory functions typical of mobile genetic elements, such as integrons (72), restriction-modification (R-M, (73)), toxin-antitoxin (TA, (74)), and exclusion (42) systems. We identified two integrons in ICEs of the SXT family (MPF_F_) (as first shown in (75)) and one array of *attC* sites lacking the integron-integrase (CALIN elements (37)) in another ICE. We identified 23 ICEs with at least one R-M system (all four types of R-M systems could be identified). The frequency of R-M systems in ICE (12% of all elements encoded at least one complete system) is similar to that observed in plasmids (10.5%) and higher than in phages (1%). Interestingly, as it was the case in these MGEs (73), the frequency of solitary methylases (25%) was higher than that of the complete systems. These solitary methylases might provide a broad protection from the host R-M systems. Most RNA genes identified in ICE corresponded to intron group II associated RNAs (36% excluding type G), but MPF_G_ ICEs encoded many *radC* and STAXI RNA (80% RNA in MPF_G_). Both genes are associated with anti-restriction functions (76). They might defend the element from R-M systems, thus explaining the relative rarity of the latter in this family of ICEs (7.5% vs 11%). Few ICEs encoded entry exclusion systems (6%, mostly in MPF_T_). One ICE contained a type II CRISPR-Cas system in *Legionella pneumophila* str. Paris. Overall, genes encoding many molecular systems associated with plasmid biology could be identified in ICEs, even if some were relatively rare.

### The organization of ICE

We grouped ICEs by their MPF types and analyzed their genetic organization (Figure 6 and Figure S11). We restricted our attention to ICEs with one single integrase of the Serine or Tyrosine recombinase families (141/199), to avoid the inclusion of recombinases with functions unrelated to the integration of the element and to facilitate the representation of the ICE organization. We represented ICEs in such a way that the integrase was located in the first half of the element. Actually, almost 90% of the Tyrosine and Serine recombinases were within the five first percent of the ICE, as expected given their role in the integration of the element. DDE transposases were randomly distributed within the ICE (Kolmogorov-Smirnov test, p-value=0.27), which suggests that most of them are not involved in the integration of the ICE. The transposases in the inner parts of the element may be involved in accretion and deletion of parts of the element, and can lead to its integration in the chromosome in regions that would not be targeted by the other integrases.

**Figure 6:**
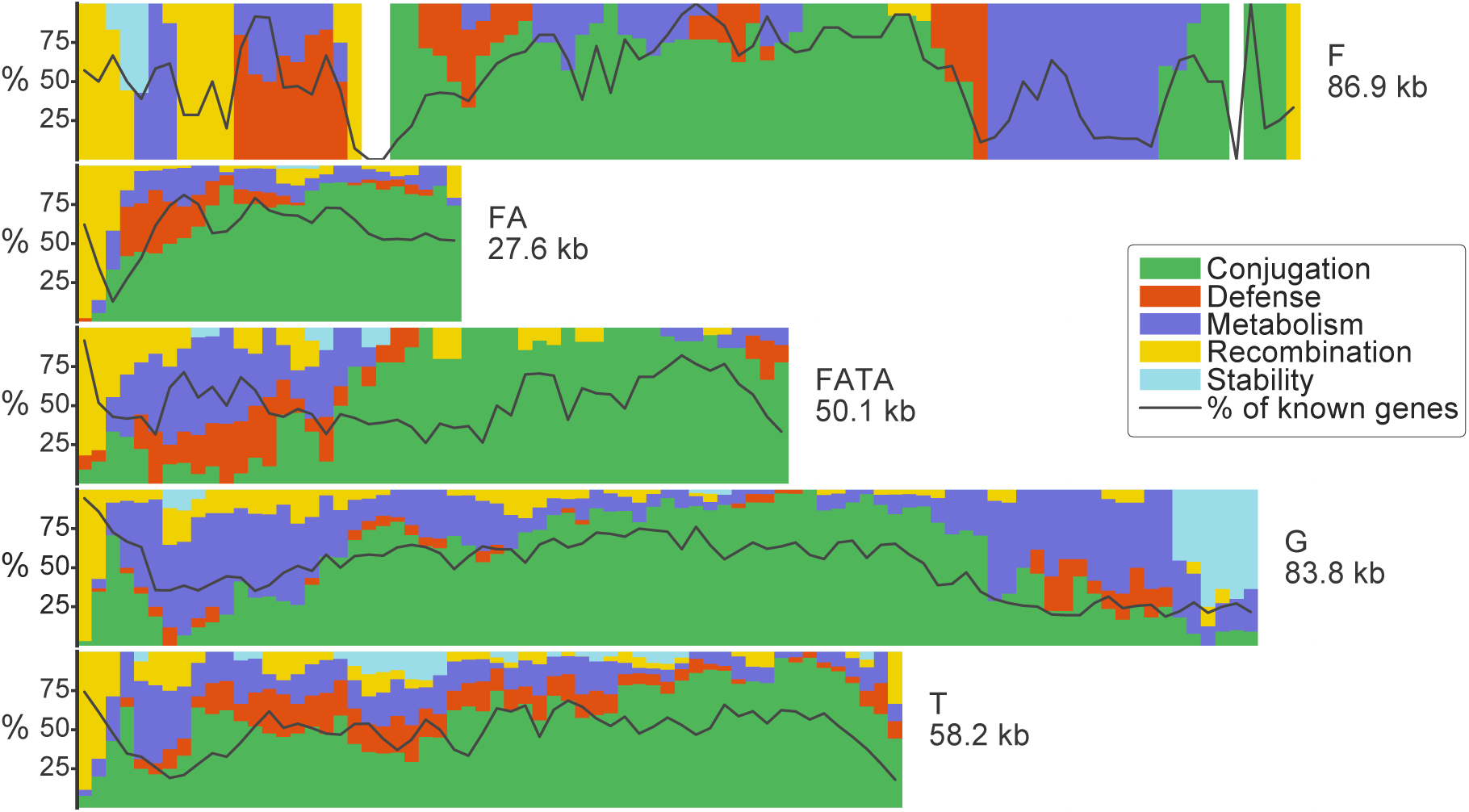
Average organization of ICEs. Each row represents an MPF type and its length is proportional to the mean size of the ICEs of the corresponding type. Colors represent different classes of functions. The black line represents the proportion of genes of known function per bin. The classes of functions correspond to Figure S9-A-B-C-F. More precisely, Conjugation includes MPF-associated genes, and the relaxase. Defense includes antibiotic resistance genes, restriction modification, solitary methylases. Metabolism includes genes annotated by EggNOG as such. Recombination includes tyrosine and serine recombinases, and DDE transposases. Stability includes replication, partition and entry exclusion systems. Bar heights are proportional to the proportion of genes of a given function at that position among the genes of known function. The width of each bin is 1kb.

To facilitate the representation of the organization of ICE, we ordered the gene orientation relative to *virB4*, which was placed on the top strand. Expectedly, given that they are often part of the same operon, the remaining components of the T4SS were usually found in a single locus and almost always (>96%) on the same strand as *virB4* (Figure S11A). The T4SS locus spanned, on average, 26% of the ICEs (Figure 6 and S12). As observed in plasmids (7), the relaxase (MOB) gene was sometimes encoded close to the T4SS genes (MPF_F_, MPF_T_, MPF_FA_) and sometimes apart (MPF_G_, MPF_FATA_). Interestingly, the relaxase and *virB4* were encoded in the same strand in most cases (86%).

Most accessory functions were encoded apart from the T4SS genes, with the known exception of the entry exclusion systems (42) (Figure S11C). We showed above that partition and replication functions co-occurred in ICEs. Here, we show that they colocalize within the element. In MPF_G_, they are often found in the edge opposite to the integrase, whereas they are close to the integrase in MPF_F_. The genes classed as “Metabolism” were also encoded away from the region of the T4SS, and were typically found in discrete modules. Intriguingly, some regions (*e.g.*, integrase-proximal regions in all types and also at the opposite end in MPF_G_ and MPF_F_, Figure 6) were particularly rich in genes of unknown function. The MPF_T_ ICEs were an exception to most of these trends, since their genes were almost uniformly distributed along the elements. This group may be genetically more diverse than the others, which would explain these results and the scattering of these ICEs in the homology network.

Interestingly, almost all genes in the ICEs were encoded in the strand of *virB4* (>80%), including the RNA genes (Figure S11F). Furthermore, genes were predominantly encoded in the leading strand for all types of ICEs but MPF_G_ (Figure S13). Overall, these results show a certain level of modularity in the organization of ICEs, as previously described in phages and plasmids (77,78), and frequent co-orientation of genes, as identified in different ICE families (SXT (79), Tn916 (80)) and in lambdoid phages (81).

### The chromosomal context of ICE

We analyzed the chromosomal context of ICEs to characterize their integration patterns. The analysis of the two chromosomal genes bordering the elements showed that ICEs are often integrated near hypothetical proteins (52%) or tRNAs (30%). These tRNAs decoded 12 different amino acids, in most cases Leucine, Lysine, and Glycine. The tropism towards integration near a tRNA varied with type of ICE, it was high for MPF_G_ (75%) and null for MPF_FATA_ (0%) (Figure 7).

**Figure 7:**
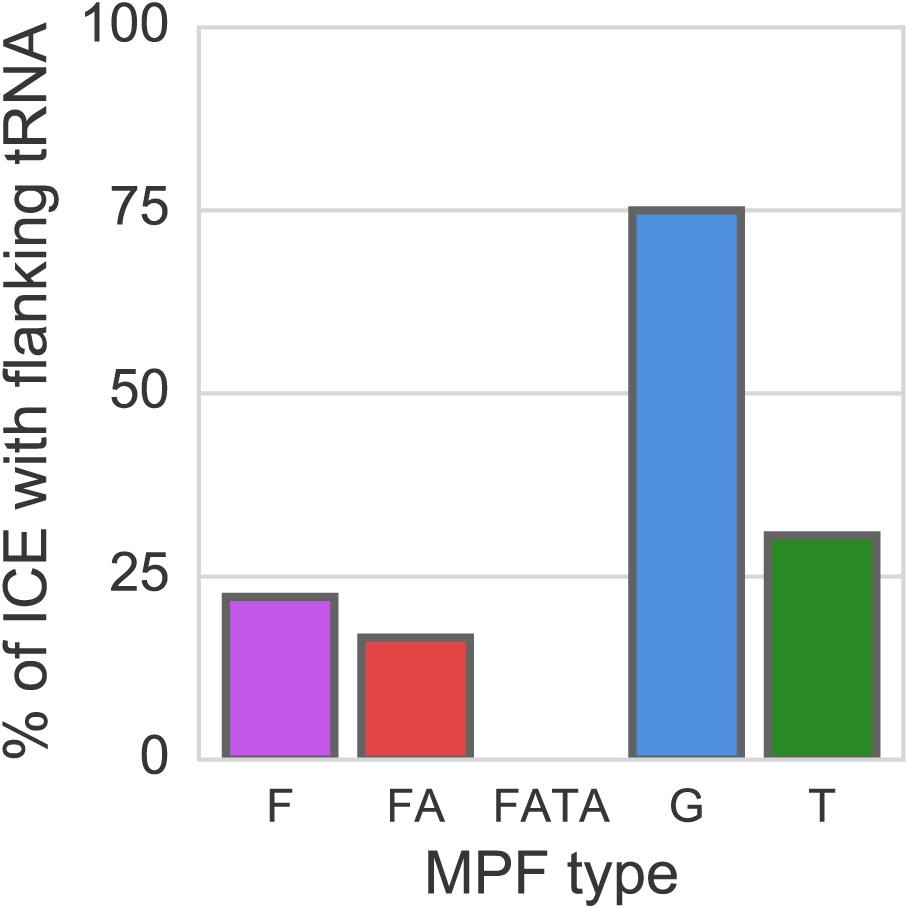
Proportion of ICEs with flanking tRNAs on either side of the ICE.

We then analyzed the distribution of ICEs in the larger context of the bacterial chromosome. Based on the analysis of 15 ICEs, it had recently been suggested that ICEs would be more frequent close to the origin of replication because they target essential highly conserved genes (26), which are more frequent near the origin of replication in fast growing bacteria (82). However, we could not find a significant correlation between the frequency of ICEs and their position in the origin-to-terminus axis of replication (Figure 8). When we analyzed the chromosomal distribution of ICEs across MPF types, MPF_FATA_ were over-abundant in the terminus region (χ^2^, p-value<0.003), whereas the others didn’t show significant trends (p-value>0.1, same test). Neither the strand location (χ^2^, p-value=0.39) nor the size of the ICE (Spearman-ρ, p-value=0.52) were associated with its distance to the origin of replication.

**Figure 8:**
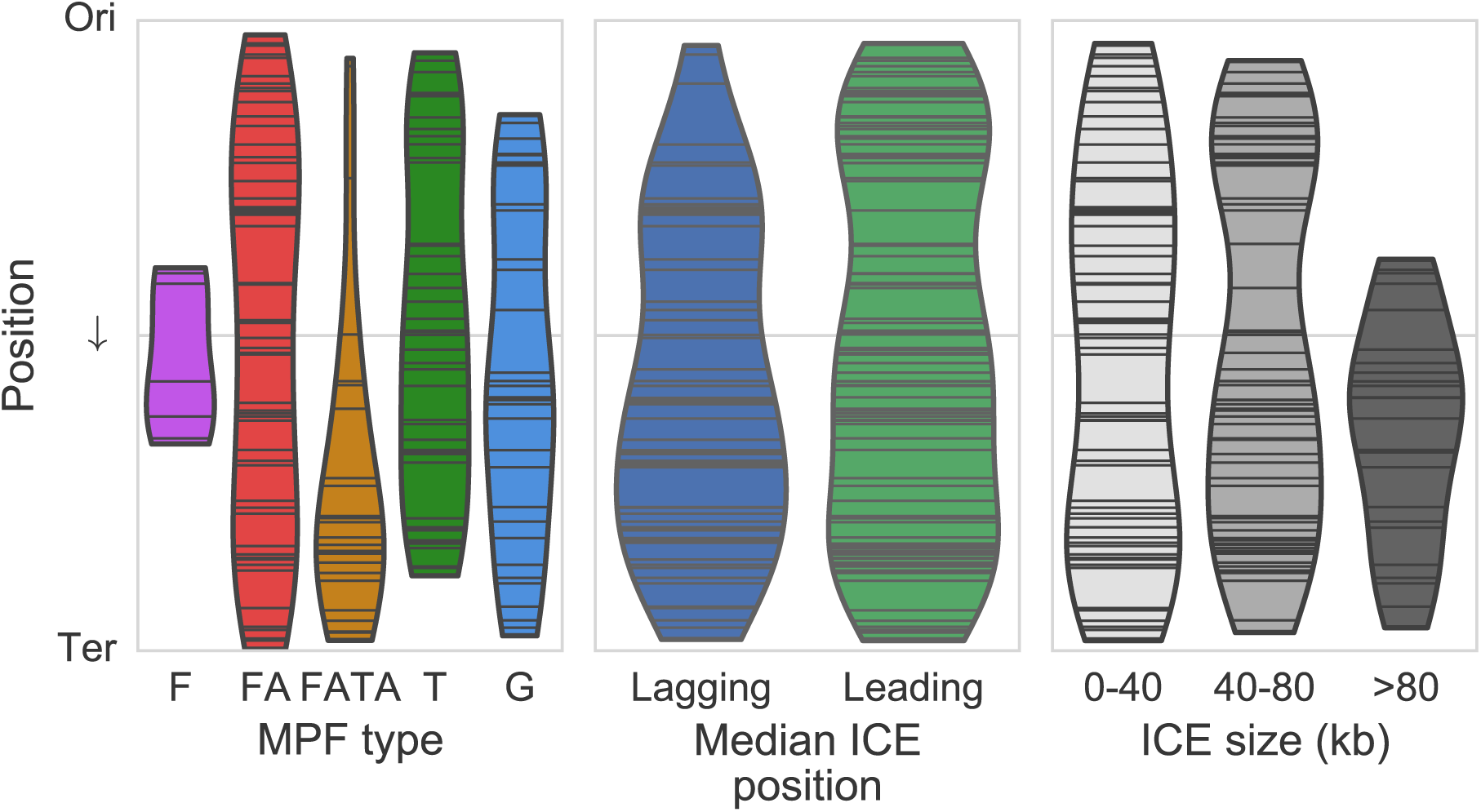
Position of ICEs along the Ori-Ter axis according to their MPF type. Left: distribution of the ICE locations as a function of the MPF type. Only type FATA is more frequent towards the half closer to the Ter region (χ^2^ test, p-value<0.001). Center: distribution of ICE locations as a function of its median strand position. Right: distribution of ICE locations as a function of the size category of the ICE.

We identified the origins and terminus of replication of genomes and inferred the leading and lagging strand of each gene in ICEs (see Methods). Most genes were oriented in the same direction as the replication fork (leading strand, χ^2^, p-value=10^-5^), with the exception of MPF_G_ ICEs that showed an opposite trend (Figure S13). The high frequency of leading strand MPF_FATA_ ICEs may be associated with the hosts′ genome organization, since they are all Firmicutes in this set and they are known to show high frequency of genes in the leading strand (83).

## Conclusion

Our work shows that one can identify and delimit ICEs from genome data using comparative genomics. The precise identification of integration sites and the validation of the functions of these ICEs will require further experimental work by experts on a large number of different species and conjugation systems. In this respect, a current limitation of our approach is the reliance on a set of complete genomes for a given species. This means that ICEs from poorly covered taxa could not be studied. Even if our dataset is representative of the diversity of experimentally studied ICEs, we lacked ICEs from types B (Bacteroidetes), of which there are experimental systems (84), and C (Cyanobacteria), for which there is no experimental system. The growth of genome databases, or the ability to include information on draft genomes (not yet possible in our pipeline), will soon correct this problem. Another limitation of this work is the assumption that the presence of a certain number of components of the T4SS system and a relaxase are necessary and sufficient to define an ICE. While our previous studies have shown that we are able to identify known conjugative systems accurately, we cannot exclude the possibility that some of the identified ICE are defective for transfer. This may explain the existence of some elements that lack identifiable integrases. In spite of these limitations, the availability for the first time of large dataset of ICEs having undergone a systematic expert curation allowed to quantify many traits associated with ICEs and confirm, and sometimes infirm, observations made from the small number of well-known ICE models. It also allowed to characterize their genetic organization, and identify common traits.

Integration and conjugation are the two only functions that we found to be present in most ICEs. Interestingly, even these functions had very different phylogenetic histories, as revealed by the scattered distribution of ICEs per MPF type in the phylogenetic tree of the tyrosine recombinases. A number of other functions were often identified in some types of ICE, notably defense systems, partition, and replication. Collectively, they reinforce the suggestions of a thin line separating ICEs from conjugative plasmids (26). The analysis of the protein sequence similarity networks between ICEs shows that these functions are often homologous among elements, tending to cluster by MPF type. The analysis of the genetic organization of ICEs suggests that they are organized in functional modules. Together, these results suggest that ICEs are highly modular, which may contribute to the evolution of their gene repertoires by genetic exchange between elements, as previously observed in temperate phages (77). If so, our data suggests that either ICEs tend to recombine more with elements of the same MPF type, or that the fitness of the products of recombination tends to be higher when recombination takes place within ICE of the same type (*e.g.*, for functional reasons).

## Availability

The program to identify conjugative systems is available on https://github.com/gem-pasteur/Macsyfinder_models

The webserver is hosted on: https://galaxy.pasteur.fr/

The program and data to make representations like those of Figure S1 is available at https://gitlab.pasteur.fr/gem/spot_ICE.

## Acknowledgements

J.C. is a member of the « École Doctorale Frontière du Vivant (FdV) – Programme Bettencourt »

We thank Christine Citti for insightful comments on a previous version of this manuscript, Fernando de la Cruz for insights and providing the list of rep proteins of PLACNET, Bertrand Néron and Olivia Doppelt-Azeroual for setting up the webserver version of CONJscan.

## Author contributions

JC and EPCR designed the study. JC and MT produced the data. JC made the analysis. JC and EPCR drafted the manuscript. All authors contributed to the final text of the manuscript.

## Funding

European Research Council [EVOMOBILOME, 281605 to E.P.C.R.]. Funding for open access charge: ERC EVOMOBILOME.

## Conflict of interest statement

None declared

